# Gaining Insight into the Deformation of Achilles Tendon Entheses in Mice

**DOI:** 10.1101/2021.01.23.427898

**Authors:** Julian Sartori, Sebastian Köhring, Stefan Bruns, Julian Moosmann, Jörg U. Hammel

## Abstract

Understanding the biomechanics of tendon entheses is fundamental for surgical repair and tissue engineering, but also relevant in biomimetics and palaeontology. 3D imaging is becoming increasingly important in the examination of soft tissue deformation. But entheses are particularly difficult objects for micro-computed tomography because they exhibit extreme differences in X-ray attenuation. In this article, the ex vivo examination of Achilles tendon entheses from mice using a combination of tensile tests and synchrotron radiation-based micro-computed tomography is reported. Two groups of specimens with different water content are compared with regard to strains and volume changes in the more proximal free tendon and the distal tendon that wraps around the *Tuber calcanei*. Tomograms of relaxed and deformed entheses are recorded with propagation-based phase contrast. The tissue structure is rendered in sufficient detail to enable manual tracking of patterns along the tendon, as well as 3D optical flow analysis in a suitable pair of tomograms. High water content is found to increase strain and to change the strain distribution among proximal and distal tendon. In both groups, the volume changes are higher in the distal than in the proximal tendon. These results support the existence of a compliant zone near the insertion. They also show that the humidity of the specimen environment has to be controlled. Necessary steps to extend the automatic tracking of tissue displacements to all force steps are discussed.

## Introduction

In the locomotor system, bones mainly provide bending stiffness,^[1]^ while tendons and ligaments transfer tensile loads.^[2]^ These load cases merge at insertions of tendons and ligaments into bone, called entheses. Insertions to the ends of long bones usually share a characteristic transition in tissue composition, structure and properties, termed the fibrocartilaginous enthesis.^[3]–[8]^ The collagen fibers pass through unmineralized and mineralized fibrocartilage before they separate into fibrils that seem to be continuous with the collagen fibrils of bone.^[9]^

### The mechanical problem of hard-soft transitions

Transitions between hard and soft materials in engineering are prone to the occurrence of stress maxima.^[10]–[12]^ If tension is for example transferred normal to the interface, the hard material prevents the neighboring soft material from transversal contraction. Thereby, it attracts stress and experiences a locally increased equivalent stress. To date, it is unclear whether and how the enthesis achieves a homogeneous distribution of stress along and across the transition from soft to hard tissues. Unmineralized and mineralized fibrocartilage are thought to develop in response to the specific loading conditions at the enthesis.^[3][6][13]–[17]^ Understanding how the enthesis leads to homogeneous stress at the interface between the different stress regimes of tendon and bone and how it mediates their different modes of deformation, is fundamental for the development of techniques for surgery and tissue engineering. It could also promote the deduction of loading history from fossil bones and inspire engineers to develop more robust connections between textile constructs and rigid frameworks. In order to understand enthesis biomechanics, macroscopic deformations have to be related to the underlying microscopic deformations.

### Biomechanical examinations of tendon-bone insertions

There is a long record of examinations into the tensile behavior of tendons and ligaments. The diversity of methods used and structures examined is vast. We reviewed some studies that include a comparison of the midsubstance and the insertion and give a gross summary. Commonly, higher strains and lower moduli are found near the insertion than in the midsubstance. Only a few studies in intact tendons find the highest strains in or towards the tendon midsubstance.^[18][19]^ Several studies in intact tendons^[20]–[22]^ find strains at the insertion to be about twice as high than in the midsubstance – one found a higher difference.^[23]^ Examinations in which the specimens were cut generally tend to find more pronounced differences between a stiff tendon or ligament and a compliant insertion.^[5][8][24]–[26]^ Only some studies of the biomechanics of tendon-bone insertions are based on a high local resolution: By a combination of X-ray dispersive spectroscopy and tensile tests in microscopic beams, Deymier et al.^[7]^ were able to identify a region within the mineralization gradient that is more compliant than the neighboring unmineralized fibrocartilage. Four other studies also confirmed the existence of a very localized compliant zone.^[8][25][27][28]^

### Unresolved issues of enthesis mechanics

This study mainly focuses on the qualitative analysis of enthesis mechanics. But a thorough pursuit of the presented approach including a quantitative full-volume analysis of the deformation might be able to resolve the following questions:

1. The reviewed publications suggest that a zone with a higher compliance than either bone or tendon is present at many insertions. Recent findings and models support the notion that tendons and fibrocartilages are poroelastic.^[29]–[31]^ Slicing significantly alters tissue biomechanics by generating an open surface where pressurized liquid can evade.^[32]^ Therefore, it is still not clear, whether the high strains at the insertion have to be partly attributed to the destruction of tissue structure by slicing.
2. A narrow region with high compliance could well be the reason for the broader finding of a compliant insertion. To check this, the local distribution of tissue strain along the transition from tendon to bone has to be investigated in intact specimens at high resolutions.
3. Connizzo and Grodzinsky^[30][31]^ examined tendon surfaces by nanoindentation. They found a poroelastic response with a higher tissue permeability near the insertion than in the free tendon. These findings are in contrast with adaptionist models of the biomechanics of wrap-around tendons^[33]^ suggesting that a lower permeability in compressed regions and pressurization of the liquid phase could reduce stress in the solid phase. In the nanoindentation experiments by Connizzo and Grodzinsky,^[30]^ the adjacent bone surface was removed from the tendon, so that the tissue did not experience physiological compression. Volume changes in the tendon and liquid accumulation in its surrounding, e.g. in neighboring bursae, should be measured to provide reference values.
4. Sevick et al.^[32]^ reported that fibers change their angle over a very short distance at the insertion into hard tissue under load. They suggest that the fiber curvature radius should not fall below a minimum determined by the bending strength of the fibers. However, increased equivalent stress resulting from the pressure exerted by the neighboring matrix could as well determine the allowable minimum curvature.^[34]^ Because of the poroelasticity of the tissue, the angular changes require reexamination in intact specimens in which liquid cannot be pressed out at the surface.
5. Several studies raised the question, whether there are mechanisms that could lead to a homogeneous loading of the enthesis in spite of varying insertion angles.^[8][18]^ Testing devices should be designed in a way enabling changes in the angle between tendon and bone in a physiological range to search for such mechanisms.

### Synchrotron radiation and the analysis of soft tissue deformation

The research questions on enthesis biomechanics demonstrate that future examinations should be carried out in intact specimens. Furthermore, full-volume approaches are required to avoid a surface bias and to account for the “subsurface nature of interfacial tissue”.^[35]^ Magnetic resonance imaging (MRI) was applied to examine intact specimens^[18]^ and to even measure strain in the full volume.^[23]^ But fibrocartilages are very heterogeneous at a scale below the resolution of MRI.^[8][36]^ The tensile response of tendon is determined by microscopic deformation mechanisms like fiber realignment and sliding.^[21][37]^ An imaging technique that can resolve the fibers and visualize the complete specimens at the same time could help to relate macroscopic biomechanics of tendon-bone insertions to the deformation of microscopic structure. Accordingly, a resolution of few micrometers is required within a field of view spanning several millimeters for a murine tendon.

Synchrotron radiation-based micro-computed tomography (SRμCT) combined with automatic tracking of tissue displacements has the potential to provide such a link between macroscopic tissue mechanics and micromechanics.^[38][39]^ Large sensors with a high pixel density provide the required combination of a high resolution and a large field of view. The coherence of synchrotron radiation allows for the visualization of untreated soft tissue samples using grating- or propagation-based phase contrast as exemplified by recent studies.^[39]–[41]^ Avoiding contrast agents is important because they are likely to alter the mechanical properties of the tissues.^[42][43]^ Due to high photon fluxes, specimens can be scanned at high frame rates and within short tomography times. High temporal resolutions are required to approximate tendon viscoelastic and poroelastic behavior. Even tomography times in the order of ten seconds appear long when compared with in vivo dynamics. Short tomography times are as well a means for keeping the radiation dose low.

Tendon-bone insertions are difficult specimens with regard to soft tissue contrast. The mineral content of bone and mineralized fibrocartilage brings about high X-ray attenuation, while the X-ray attenuation of tendon and unmineralized fibrocartilage is only slightly above that of water. Here, we explore a workflow and parameters for propagation-based phase contrast imaging. We report some exemplary results, as well as difficulties, solutions, limits and potentials of this approach. In a single specimen, we also examine which further steps are required to achieve full-volume displacement tracking over complete deformation tests.

## Results

### Forces, relaxation and rupture

Specimens withstood a force program of one to four force steps. Most often they failed when the force was increased to force level three after the third tomography. In tests with a higher target force in the first force step, failure sometimes occurred during the second force step. A control specimen that was not exposed to radiation during the tomography phase did not fail within six force steps (Fig. S1). At the last force level a target force of 1.65 N was reached.

The relaxation behavior of the specimens (Fig. 1A) can be approximated with logarithmic curves. In a watered and an unwatered specimen, the measured force decreased to 62 % of the target force within the relaxation time at force level one and 63 % respectively, and further to 54 % and 59 % of the target force within the following tomography phase. In the control specimen, the measured forces were 66 % and 58 % of the target forces at corresponding times in the force program.

**Figure 1:**
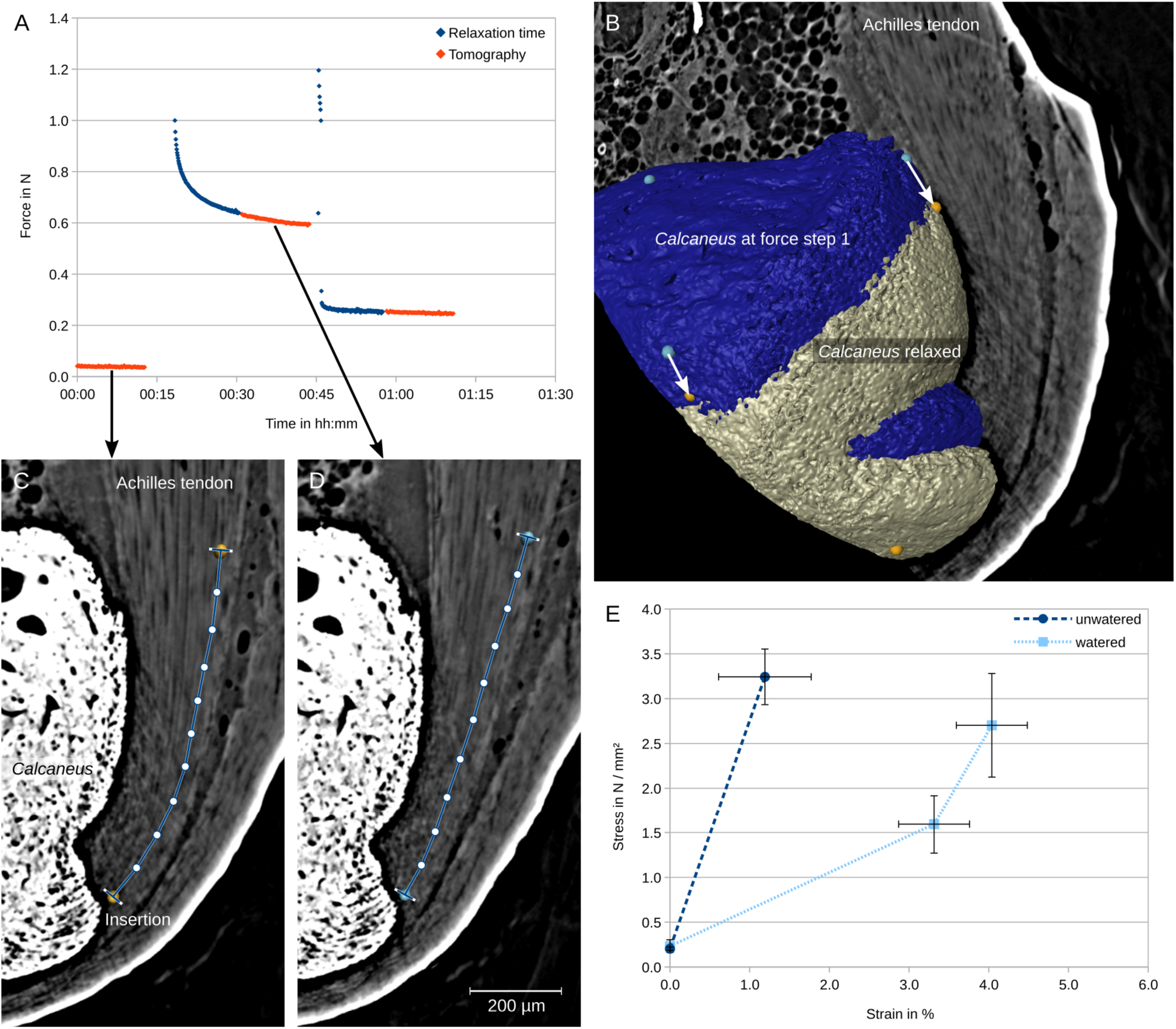
Analysis of tensile behavior. (A) Force is logged during relaxation time and tomographies. (B) Rendering of the Calcaneus surfaces of a specimen in the relaxed state (reference, beige) and at force level one (blue). The section corresponds to the reference specimen. Markers are assigned to corresponding surface patterns and used for the registration of the volume image from force level one to the reference as illustrated by arrows. (C) Co-registered sections through an unwatered specimen in relaxed state and at (D) force level one with landmarks and semilandmarks for strain measurement. (E) Diagram of stress over strain in unwatered and watered specimens (mean values with standard deviation, n=3).

### Imaging and deformation of the soft tissues

Propagation-based phase-contrast imaging renders the hard tissues with surrounding phase-contrast fringes (Fig. 1B–D). The *Calcaneus* is the largest object with a strong attenuation in each specimen. Furthermore, in many specimens mineralizations are found in the free tendon at the proximal end of the dataset. In regions where the beam paths passed through hard tissues, the 3D reconstructed image is noisy. Especially adjacent to the *Calcaneus*, near the insertion, artifacts are superimposed on the tendon structure. However, in many regions of the tendon, the fibers can be identified. The gray values in tendon tissue are higher than in the surrounding soft tissues. The bursa between the *Calcaneus* and the tendon is identified by the homogeneous gray value of the contained liquid. The fat pad that is situated anterior of the tendon and proximal of the bursa exhibits a characteristic cellular structure.

The patterns that were tracked by human subjects correspond to inhomogeneities along the tendon fibers like cellular lacunae, varying gray values, branching patterns and characteristic arrangements of fibers and non-fibrillar components. At position P0 near the insertion the distinction of such patterns from the superimposed artifacts is difficult. At the same time, the displacements in the coregistered datasets are small near the hard tissue interface. The volume rendering at position P1 as well exhibits many artifacts. At position P2 the displacements are large. In some specimens, tendon mineralizations are useful as markers or for a validation of nearby patterns.

The deformations during the first force step are larger than during the second one. The orientations of the fibers clearly increase during this force step. Proximal of P1 a region with macroscopic fiber kinks is seen in two watered specimens. The kinks are straightened during the first force step.

The angles between fibers and the mineralization front become smaller with increasing load at an angle of load application of 90° (Fig. 1C–D, 2A–C), but not at an angle of 120°. A sharp angular change in the fiber direction is observed within a narrow band near the hard tissue surface.

**Figure 2:**
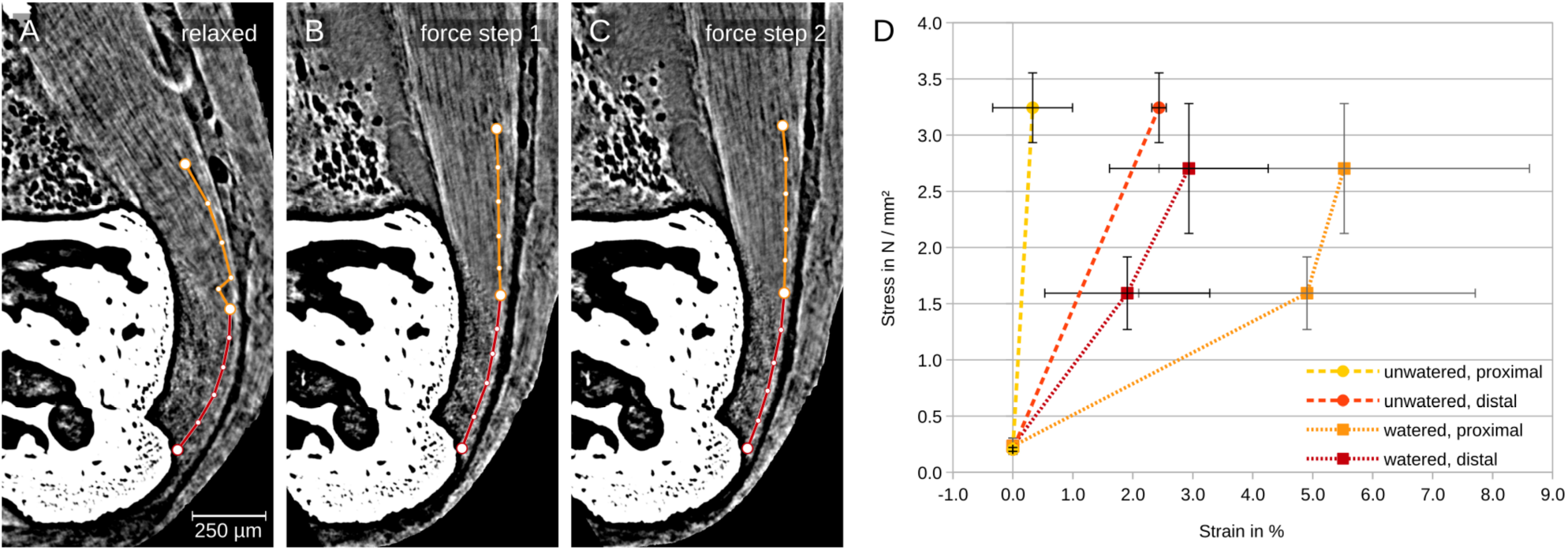
Tensile behavior of proximal and distal tendon segments. (A) Co-registered sections through a watered specimen in relaxed state and at (B) force level one and (C) force level two with landmarks and semilandmarks for the proximal (orange) and distal (red) measuring distance. (B) Diagram of stress over strain in the proximal and distal measuring distance in unwatered and watered specimens (mean values with standard deviation, n=3).

### Tensile data

The tendon cross-sectional area (CSA) at position P1 was 0.218 ± 0.029 mm^2^ in watered specimens and 0.177 ± 0.014 mm2 in unwatered specimens. The difference is not significant.

The mean force during the tomography and the resulting stress at P1 differ widely between the specimens within each of the force levels.

In all cases, the first force step still comprises a part of the toe region of the stress-strain curve. The mean tangent modulus of the first force step (42 ± 15 MPa) is significantly lower than that of the second force step (151 ± 55 MPa) in the watered specimens (Fig. 1E, S2). In the single unwatered specimen, that lasted up to the end of the tomography at the second force level, the tangent modulus as well increases from the first (356 MPa) to the second force step (722 MPa). In unwatered specimens, the mean modulus in the first force step (287 ± 97 MPa, Fig. 1E) is significantly higher than in watered specimens. In the single specimen tested at an angle of load application of 120° the tangent modulus in the first force step is similar to that of the other unwatered specimens (Fig. S2). In the second force step, no increase in strain was measured in spite of an increase in stress.

### Differences between distal end and free tendon

The regional strain measurements along the proximal and the distal tendon partly yielded negative strain values (Fig. S3, S4). Such findings occurred in watered and unwatered specimens and at both force levels. We therefore refrain from quantifying the moduli for these regions and just give a qualitative description.

The following descriptions refer to the first force step. In unwatered specimens, significantly higher strains were measured in the distal region as compared to the proximal region. In watered specimens, the relations among proximal and distal region tend to be inverse (Fig. 2D, S3, S4). However, there are no statistically significant differences between the strains and moduli. The modulus of the proximal region is relatively low. The moduli measured in the distal region of watered and unwatered specimens are very similar.

The regional modulus values exhibit a significantly higher variation in the watered specimens than in the unwatered specimens. The variation is even high between samples that were subjected to similar levels of stress (Figs. S3, S4). In the watered specimens, the moduli of both regions seem to increase in the second force step.

### Volume losses

The mean relative volume loss in the distal tendon seems to be higher than in the proximal tendon irrespective of water swelling (Fig. 3). In the watered specimens, the relative loss during the second force step tends to be smaller than the initial volume loss (Fig. S5). In the proximal as in the distal tendon, the volume further decreases by 2–5 % in the second force step with respect to the volume in the relaxed state. In the unwatered specimens, the sample size in the second force step is too low to allow statements about volume losses (Fig. S6).

**Figure 3:**
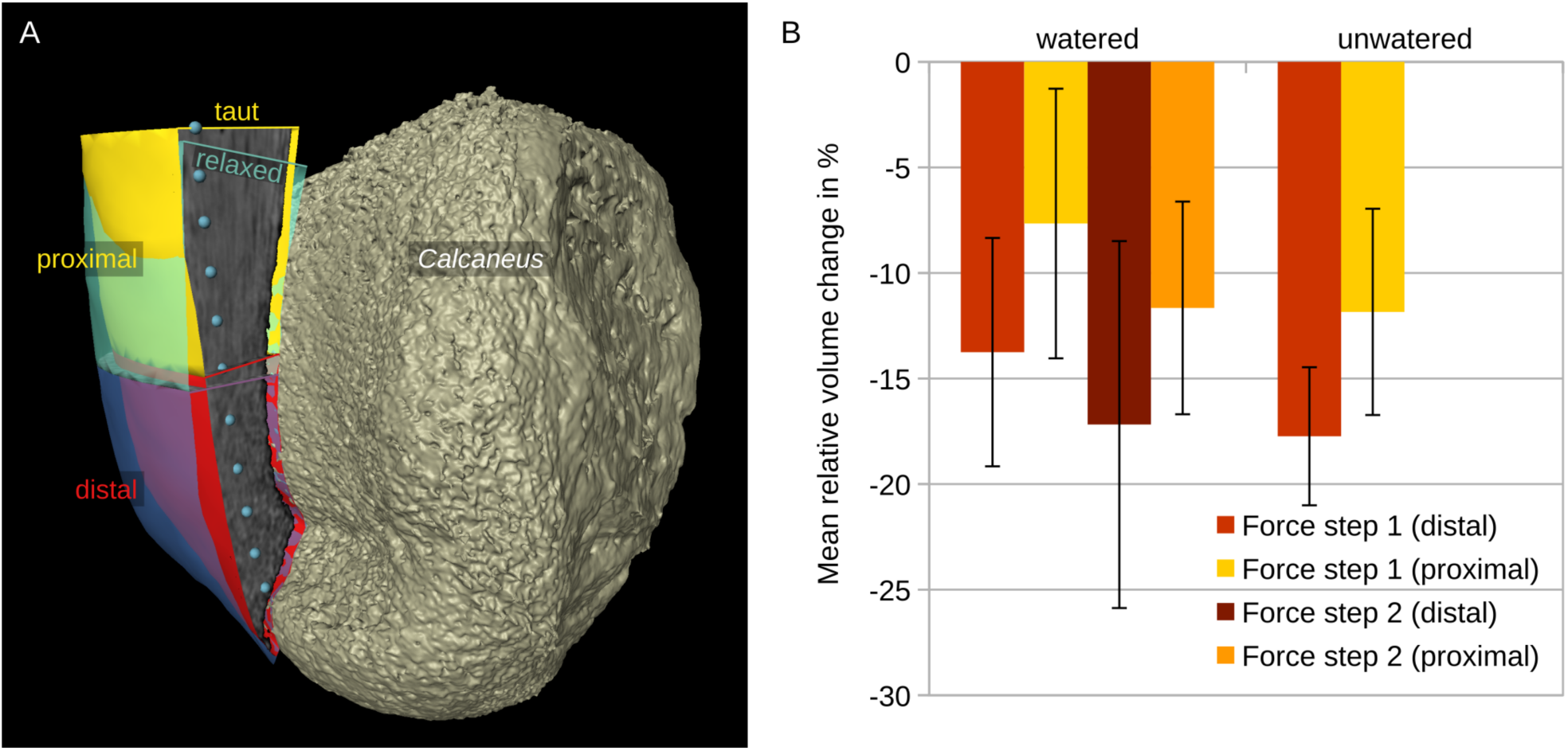
Volume changes due to stretch. (A) Rendering of the surfaces of the proximal and the distal segment of an unwatered specimen in relaxed (cyan and purple) and stretched state (yellow and red). (B) Diagram showing the volume changes in the proximal and the distal tendon segment with respect to the volume in the relaxed state in watered and unwatered specimens (mean values with standard deviation, n=3).

In the proximal tendon, there is an interesting dependency of the volume loss on the examined proximal length: The relative volume loss is the higher, the further the examined volume extends in the proximal direction (Fig. S7). The dependency is found in the watered specimens in both force steps and in the unwatered specimens.

In some specimens, increases in liquid volume in the bursae anterior and posterior of the tendon were observed (Fig. 2A–B). In the single specimen tested at an angle of load application of 120°, the volume loss in the tendon is lower than in the other specimens. The absolute increase in bursa volume is in the same range as the volume loss in the tendon in this specimen (Fig. S8).

### Findings from the optical flow analysis

With only low contrast available in the soft tissue, a global optimum for the displacement field was not found in the larger deformations of the first force step. But a plausible result was provided for the second force step. The Green-Lagrange strain tensors (Fig. 4) were calculated from the displacement field. The component E_yy_ of the tensor was roughly aligned with the tendon course and the direction of the tensile force. The strains in this direction were mostly positive with low absolute values. In the superficial distal tendon, negative strains were found. At P1, the component E_xx_ roughly corresponds to the radial direction of the tendon curvature around the *Calcaneus*. In this direction negative strains were found with the most extreme values in the superficial distal tendon. Negative strains with smaller absolute values were measured in mediolateral direction. This direction corresponds to component E_ZZ_. Component E_xy_ represents the shear in the sagittal plane. It shows a similar distribution as component E_xx_. The most extreme values were measured in the superficial distal tendon.

**Figure 4:**
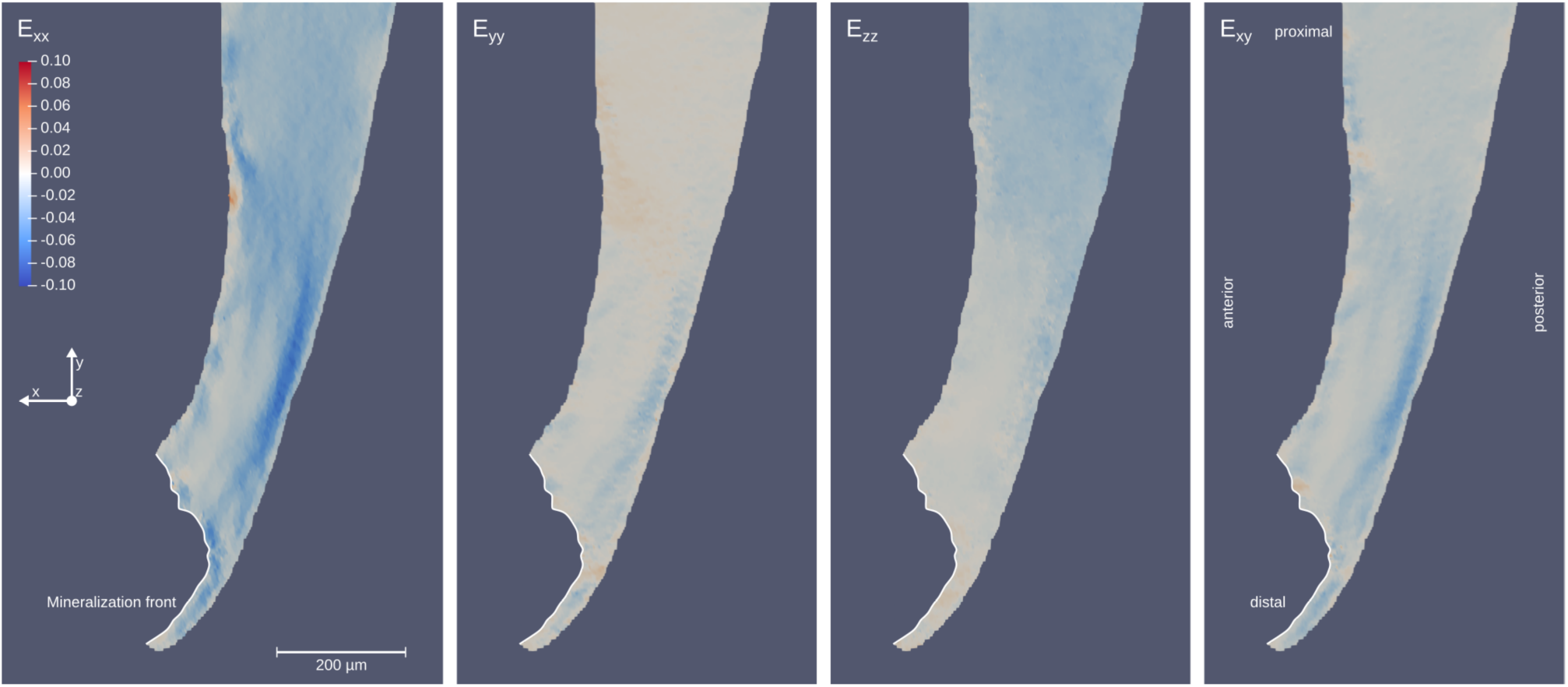
Strain fields for the second force step measured by optical flow analysis in an unwatered specimen. Several components of Green-Lagrange strain tensor are plotted over the sagittal section of the tendon.

The optical flow analysis and the visual analysis were compared with regard to the deformation of the measuring distance from the visual analysis of one unwatered specimen. Qualitatively, the optical flow analysis and the visual analysis consistently lead to the measurement of very small strains over the second force step. For the distal region the measured strains are even negative with both methods. With regard to absolute values the strains from the optical flow analysis tend to be smaller than the ones from visual analysis.

## Discussion

SRμCT with propagation-based phase-contrast was proven useful as a technique for the visualization of the deformation inside intact soft tissue specimens. Changes to the internal chemical environment of the specimens like staining were not required. With a faster procedure for dissection and mounting examinations could also be carried out in fresh or thawn ex vivo specimens without a need to soak them in phosphate-buffered saline (PBS). PBS is known to lead to tendon swelling.^[2]^ Tomograms were recorded within times short enough to preserve relevant material properties. The three-dimensional structure of tendon tissue was resolved in sufficient detail for pattern tracking.

### The effect of water content

The CSA values measured in unwatered specimens demonstrate that the histological treatments used in our previous studies^[44][45]^ like dehydration, demineralization and critical-point drying, as well as PTA-staining, led to shrinkage and underestimation of the CSA. At the same time, watering led to swelling in this study. Water uptake by watered specimens was neither controlled nor quantified. The tests in unwatered specimens are considered more reproducible. The water content of the specimens in future tests could be controlled by raising the relative humidity of the air within the specimen chamber to nearly 100 % using a humidifier.^[46]^

### Tensile behavior

As seen from the increasing orientation of fibers, the first force step still comprises a part of the toe region. In contrast, the second force step is expected to reflect the linear region of the stress-strain relation.

Ansorge et al. (2011) examined the free Achilles tendon in mice aged 28 d in a PBS bath and published a toe modulus of 72.3 ± 48.7 MPa, a strain of 1.91 % at the transition from toe to linear region and a linear modulus of 544.0 ± 240.8 MPa. Compared to this reference our study differs with regard to three conditions. (1) The age of the mice we examined is higher. Therefore, higher moduli and lower strains are expected to occur. (2) We used a much higher preloading, so that the stress-strain relation contains a smaller part of the toe region. (3) We did not determine the immediate relation of stress and strain, but the relation after a relaxation time. As reported, the stress after relaxation amounts to about 60 % of the stress measured at load application. Taking these considerations together, the stress-strain relation found in the unwatered specimens corresponds well to the values from the reference.

The higher age of the examined mice (C57BL/6J) also explains the occurrence of tendon mineralizations.^[47]^ We do not expect the mineralizations to add substantial changes to the tensile behavior in the more distal parts of the tendon that were subject to the examinations. Connizzo and Grodzinsky^[31]^ report that differences between the supraspinatus tendon mid-substance and the insertion with regard to permeability and modulus are present in mature mice (aged 4 months) and disappear in aged mice (18 months).

### Regional differences

The existence of a distal region that is more compliant than either tendon or bone is supported by our findings of higher strains in the distal tendon of unwatered specimens. However, the modulus values in the proximal tendon were too extreme to be plausible in unwatered specimens. Additionally, the relation depends on the water content of the tissues. In the watered specimens, the moduli of the proximal region tend to be lower over the first force step than the moduli measured in other positions.

Volume losses can be related to the Poisson’s ratio of the tissue, but they can also result from compressive stress. The relative volume losses in the distal part of the specimens tend to be higher than in the proximal tendon. This is mainly attributed to the curvature of the tendon over the *Tuber calcanei*. Accordingly, tensile stress in the tendon leads to a compression of the distal tendon against the bone surface. The notion is supported by the finding that the relative volume losses are lower in a test with a higher angle of load application and insertion angle. In some specimens the liquid volume in neighboring bursae is increased after the first force step. We expect that the liquid that is pressed out accumulates in the bursae and plan to examine the volume changes in the bursae. The compression obviously is time-dependent as the optical flow analysis rendered a high transversal compression in spite of a low strain increment during the second force step.

The dependency of the relative volume losses on the proximal extent of the examined proximal portion suggests that there is a region with minimal volume changes shortly above the *Tuber calcanei*. Further proximal the volume losses seem to be higher. This could result from the transfer of liquid along the tendon. Liquid from the region compressed to the *Tuber calcanei* might accumulate in the neighboring free tendon. The distribution of volume changes should be examined at a higher local resolution to investigate this. The results from the optical flow analysis of the second force step do not support the idea of a lower transversal compression or contraction above the *Tuber calcanei*.

The changes in volume also bring about changes in the angles of the fibers at the insertion to the hard tissues. The volume renderings confirm findings in sliced specimens by Sevick et al.,^[32]^ who observed that fibers under load change direction over a short distance. This implicates very small curvature radii – and could lead to high local stress which is in contrast with previous ideas of enthesis function.^[4][14][48]^ In an unpublished series of radiographic projections, we observed that fibers are bent like that within seconds after load application, while the changes within the further relaxation time are small. However, the strain rate in the experiments was much lower than in vivo loading rates.

The initial hypothesis was that pressure in the liquid phase would have a higher influence on the mechanical behavior in intact specimens. Instead many findings from sliced specimens were confirmed, for example the presence of a compliant region and angular changes of the fibers at the insertion to hard tissue. The specimens were imaged after relaxation to avoid motion blur during the tomography. Volume losses and liquid accumulation in the bursae demonstrate that liquid movements take place during loading and relaxation. We still assume that examinations during load application and relaxation would find effects related to the pressurization of the liquid phase.

## Perspectives

### Measuring stress and strain

The tensile test of this study had some limitations. The cone used for muscle trapping allowed slippage which was compensated by local measurements of strain. If both ends of the specimen were clamped with a known compliance, an overall clamp-to-clamp strain could be calculated. An even better recording of the loading conditions could be achieved by introducing the complete test length into the field of view.

For the evaluation, we assumed a homogeneous distribution of stress and constant CSA sizes along the tendon. But we know that these gross estimations do not reflect the condition in the Achilles tendon in mice.^[45]^ Instead, stress could be estimated using finite element analyses in future studies.

Importantly, strain was measured along the fibers in the visual analysis. However, longitudinal sliding among fibers and shear of the matrix between them is a major mechanism of tendon deformation.^[37]^ Strain along the fibers relates to mechanisms like uncrimping and sliding among fibrils. Following a simplifying assumption, these strains along fibers were used as an estimate of tissue strain and for the calculation of volume changes. It would be more meaningful to measure strain at a higher local resolution and relate strain tensors to the local orientation of fibers. Such an approach would enable the distinction of deformation mechanisms in 3D.

The differences between strain values from the visual analysis and the optical flow analysis could be caused by inaccuracies of the visual approach. A more thorough evaluation including optical flow analyses over several force steps at larger sample sizes will be required to quantify the error. Nevertheless, both approaches lead to similar qualitative findings: The strains in the second force step are low. The distal part of the tendon is shortened at the position of the measuring distance in spite of an increase in force.

### X-ray radiation dose

The application of SRμCT in deformation studies of biological tissues is still restricted by the low number of tomograms and thus force levels that can be recorded before the specimen fails. The relaxation curves before specimen failure did not differ notably between the scanned specimens and the control. Instead, the specimens seem to have failed, when the X-ray radiation dose exceeded a critical level and when at the same time the mechanical load was either maintained or further increased. Measurements of the radiation dose are critical to relate the findings to the literature.^[49]^ Both, the X-ray radiation dose and the time under mechanical load, can be reduced by a minimization of tomography time. Recent developments at the beamline used show that durations of tomograms can be decreased to 15 s. X-ray radiation dose can be further decreased by utilizing a fast x-ray shutter closing the beam path between single projections. The control experiment with a specimen that is not exposed to radiation during the tomography phases demonstrates that more than five force levels could be recorded under such conditions. Accordingly, the relation between strain and stress could be explored with smaller intervals between the force levels.

### Image quality and pattern tracking

The occasional findings of negative strains in regions with a high modulus might point to inaccuracies related to the visual approach to pattern tracking or to differences in the relaxation behavior across the tendon. The latter interpretation is supported because the optical flow analysis of a single force step found negative strains as well.

The reliability of visual pattern tracking and optical flow analysis depends on image quality. As yet, patterns that identify positions along the fibers are partly concealed by superimposed artifacts. Several parameters can contribute to an improvement of image quality. (1) The previously described reduction in the duration of tomographic data acquisition would at the same time decrease motion blur. (2) If motion blur is low, higher spatial resolutions could still increase the contrast between fibers and the non-fibrillar matrix. (3) The combination of photon energy, propagation distance and parameters for reconstruction and phase retrieval still requires further examination with regard to its effect on signal to noise ratio in the studied specimens. (4) Recent technological advancements in speckle-imaging and grating interferometry lead to an increase of resolution in grating based phase-contrast SRμCT which is known to achieve a higher dynamic contrast range^[40]^ and a reduction of artifacts in specimens with heterogeneous absorbance. Grating-based phase-contrast additionally allows absolute quantitative evaluation. However, grating-based imaging leads to higher radiation doses. (5) Image quality can also be increased by post-processing of the reconstructed images and application of tools for artifact filtering.

The application of the optical flow analysis to the second force step in one specimen yielded plausible results. A further evaluation of this approach is supported. The intervals between the force levels in this study led to large displacements of patterns between the relaxed state and the first force level and prevented the application of the optical flow analysis to the complete deformation test. Shorter tomography times will facilitate the recording of more force levels before a critical radiation dose is reached.

Automatic tracking would improve the approach of this study in several ways. (1) It leads to a higher reproducibility of the measurements. (2) Displacements can be measured at a high spatial resolution (3) over the complete region of interest. (4) Three-dimensional strain tensors can be calculated from the results.

### Time-dependent phenomena

Shorter tomography times also facilitate an onset of tomographies after a shorter relaxation time because they are more tolerant for relaxation motions within the specimens. Gupta et al. hint to the fact that “a significant fraction of any load- or strain-induced changes occur within the 1^st^ few seconds”.^[50]^ To reach subsecond temporal resolutions, tomographies should be combined with series of radiographic projections in a lateral or medial view to monitor parameters like tendon thickness or the orientation of fibers near the insertion through loading and relaxation.

## Conclusion

By a combination of a mechanical test and propagation-based phase-contrast SRμCT the three-dimensional structure of Achilles tendons in mice was visualized at several defined force levels. Two findings of studies in slices were confirmed in intact specimens. (1) The region near the insertion was found to be more compliant than the free tendon. (2) The fibers near the transition to hard tissue are bent under load with respect to their orientation in hard tissue. The change in orientation is restricted to a relatively short length near the insertion.

Tomography times were identified as a central parameter to enable automatic pattern tracking. Recent developments at the beam line facilitate faster tomographies. Therefore, the combination of techniques tested is suitable for a wider application in the three-dimensional soft tissue mechanics of small animals.

## Material and method

### Animals

The insertion of the Achilles tendon is a representative example of a fibrocartilaginous enthesis. Together with the sesamoid fibrocartilage, the periosteal cartilage, the bursa between them and Kager’s fat pad protruding into the bursa, it embodies the idea of an enthesis organ^[51]^ and was termed the “premiere enthesis” for this reason.^[52]^ At the same time, it is accessible to dissection and imaging. In this study the Achilles tendon enthesis in mice (Fig. 5A) is examined because it is small enough to fit into the field of view used for high resolutions at the SRμCT beamline. Fiber size seems to show negative allometry with body size.^[45][53]^ Consequently, a better resolution of the fibers can be achieved in small mammals.

**Figure 5:**
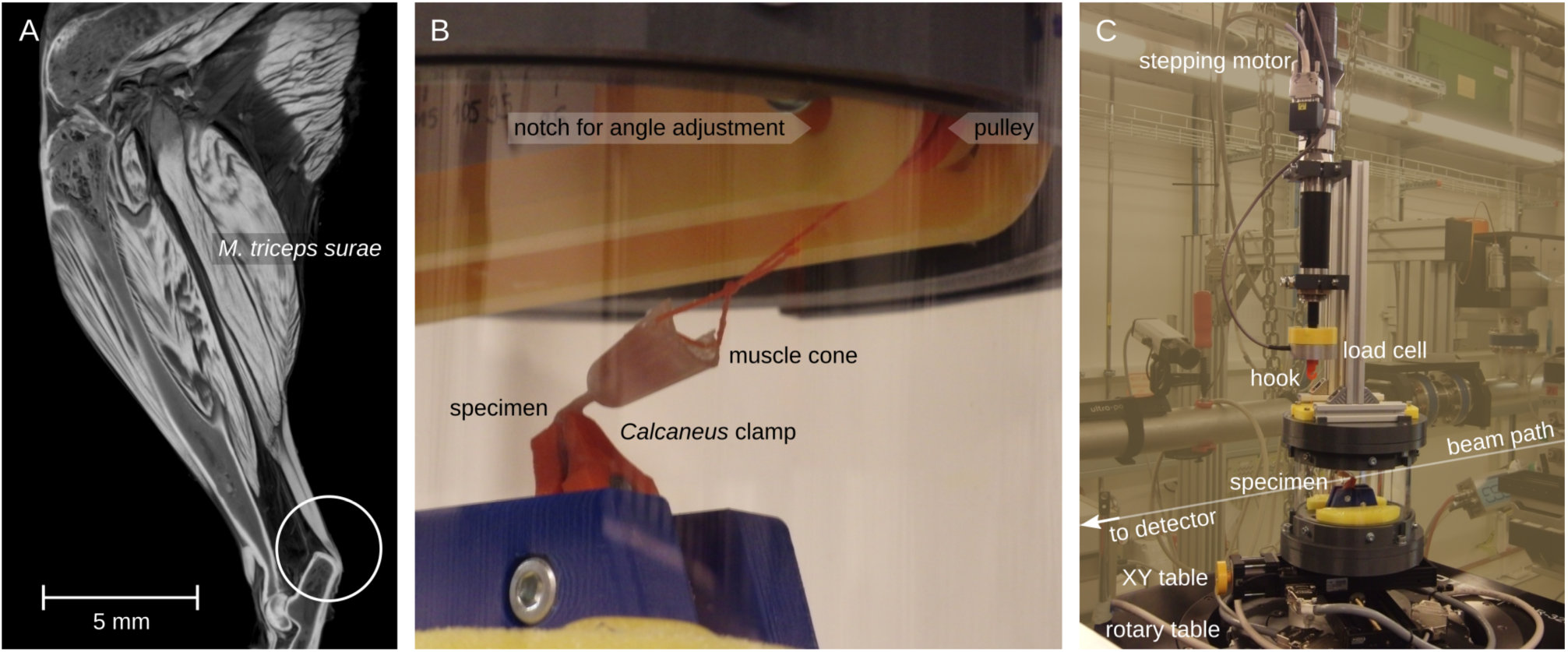
Specimen anatomy and experimental setup. (A) Volume rendering of the lower left limb of a mouse virtually cut by a sagittal section plane. The circle marks the Achilles tendon and its insertion at the Calcaneus. (B) A specimen is mounted in the tensile testing chamber, the M. triceps surae is trapped in a cone and the Calcaneus is clamped. (C) The tensile testing device allows remote control of displacement and logging of forces throughout several synchrotron radiation micro-computed tomographies. The region of interest is positioned over the rotation axis by an XY table.

Hind limbs of mice from two strains (C57BL/6J, NHE1-SMMHC | 0) were obtained freshly within 15 min after euthanasia at Institut für Allgemeine Zoologie und Tierphysiologie of the Friedrich Schiller University Jena, at the Service Unit for small rodents and the Animal Facility of the University Hospital Jena. The hind limbs were cryofixed either in isopentane cooled with liquid nitrogen or directly in liquid nitrogen. Seven hindlimbs were evaluated and are reported in this article, one further hindlimb is used as a control for the influence of radiation. The hindlimbs stem from 7 individuals from both sexes, aged 8 to 16 months that weighed between 30 g and 37 g. All procedures involving animals conformed to national and international ethical standards.

### Tensile testing device

For imaging specimens under load a custom testing device was built (Fig. 5B,C). In the path of the synchrotron X-ray beam a non-metal construction was required which was widely realized by 3D printing. An acrylic glass cylinder forms the walls of the specimen chamber. A clamp for the *Calcaneus* can be fixed to a holder at the base of the specimen chamber. A thread connects a hollow cone for trapping the *M. triceps surae* to a load cell above the specimen chamber (type 8523-5020 by Burster Präzisionsmesstechnik GmbH & Co. KG, Gernsbach, Germany) with a nominal force of 20 N. The thread runs over a pulley that is mounted on a ball bearing to minimize rotational friction. The angle between tendon and *Calcaneus* axis can be adjusted by relocating the axis of the ball bearing along a notch (“angle of load application”). This also implies a change in the insertion angle. Above the pulley, the thread forms a loop that is slipped over a hook at the load cell. The upper side of the load cell is connected to a precise linear actuator (type M-35 by Physik Instrumente GmbH & Co. KG, Karlsruhe, Germany), that can displace it with a step size of below 1 micron. Load cell and actuator are integrated in a control loop^[40]^ so that a preset force can be approached automatically. The whole device is mounted on an XY table with two linear degrees of freedom (DOFs) for horizontal positioning. The XY table is itself mounted on the sample stage of a synchrotron beamline (P05, Petra III, DESY, Hamburg, Germany) operated by Helmholtz-Zentrum Geesthacht^[54][55]^ with one rotational DOF for scanning and one linear DOF in z-direction for vertical positioning.

### Dissection and mounting

The specimens were transferred to the imaging facility on dry ice. One specimen at a time was thawed in PBS freshly taken from a fridge, and subsequently dissected. The *M. triceps surae* was separated from the shank and the knee. The plantaris tendon was isolated from the Achilles tendon and cut at mid-length. Keeping the Achilles tendon intact, the foot was separated from the shank at the ankle and placed in the clamp of the tensile testing device with its dorsal side pointing to the upper part and the posterior tip of the *Calcaneus* protruding from the tip of the clamp. The *M. triceps surae* was trapped in the hollow cone. A slit in the cone reaches to its tip. The Achilles tendon was introduced to the slit and gently pulled to the tip of the cone. Before mounting them into the tensile testing device, the clamp and the cone with the specimen were transferred to PBS again. We refrained from a continuous hydration of the specimen in a PBS bath because the higher attenuation would have decreased the contrast. The angle of load application was set to 90°. Only in the last experiment was the specimen tested at an angle of load application of 120°. The factual angle of load application can deviate slightly from these values depending on the position of the *Calcaneus* along the clamp.

As the specimen chamber is nearly closed, the humidity in the chamber can be increased to reduce evaporation. Therefore, the floor of the specimen chamber was covered with foam and paper material soaked with deionized water. In the initial experiments the specimens were rinsed with deionized water to compensate for drying during the mounting process (“watered specimens”). But this seemed to lead to swelling of the specimens. Therefore, this step was left out in late experiments (“unwatered specimens”).

### Mechanical tests

In vivo measurements^[56]^ suggest that the forces transferred by the Achilles tendon can exceed 3 N. In locomotion, forces occur dynamically and cyclically. The objective of the experiments was to examine the relation of stress and strain along the fibers in the free tendon and near the insertion. Because of the necessity to minimize motion within the specimen during the tomography, we had to examine each specimen in a quasi-static state after a relaxation phase. Measuring the load after relaxation is a common approach to the tensile behavior of tendons and their insertions.^[8][31][57]^ The tissue cannot withstand forces as high as the in vivo forces in such a quasi-static experiment because the loads have to be applied for a longer period.^[21]^

Our main input parameters in the quasi-static experiments were the target force and the given relaxation time. However, stress does not only depend on the input force, but also on the tendon CSA and the relaxation behavior in each individual specimen. Accordingly, the stresses vary between the specimens in spite of representing the “independent variable”.

In each step of the force program, the stepping motor displacing the muscle cone was actuated until the target force value was measured by the load cell. The first tomography (“relaxed state”) was recorded at a preloading of 0.05 N. In the initial experiments, various values for the next force level were explored. In the unwatered specimens, that were thought to reflect the physiological state more accurately, the target force was consistently set to 1.0 N. After reaching the target force, the displacement was maintained and the forces were logged over a relaxation time of 7–12 minutes. Afterwards, the second tomography was recorded (“force level one”). The stepping motor was actuated again to increase the force by a defined interval of either 0.2 N or 0.4 N above the first target force. After relaxation, a further tomogram was recorded (“force level two”). This protocol was followed up to the failure of the specimen.

To control for the influence of radiation, the mechanical behavior is compared to a specimen that underwent the same force program but was not exposed to radiation during the tomography phase.

### Imaging

The tensile testing device was mounted on the sample stage of the synchrotron beamline. Before the start of the mechanical testing program, the specimen was centered about the axis of rotation via the XY table. The height position of the sample stage was adjusted to place the insertion zone of the tendon at the lower end of the field of view.

The tomograms were recorded at a photon energy of 35 keV to decrease absorbance in the specimens. By a higher photon energy, the overall radiation dose a specimen is subjected to is reduced due to a reduction in absorbance. Propagation-based phase contrast makes use of the generation of intensity contrast during the propagation of the transmitted wave front. The distance between specimen and detector is thus referred to as propagation distance. It was set to 0.80 m. The optical magnification between the scintillator and the CMOS-sensor was 9.96. The CMOS-sensor comprised a grid of 5120 times 3840 pixels with a linear pixel size of 6.40 μm. The resulting field of view spanned 3.29 mm times 2.47 mm. The effective voxel size in the reconstructed tomographic volume was 1.29 μm after a twofold binning of the original projection images. For a tomography in standard mode 1,201 to 1,801 projections were recorded over an angular range of 180°. The number of projections was set to 1,800 for the same angular range in fly-scan mode with a continuous rotation of the specimen. A tomography in standard mode took between 15 and 26 minutes, while the tomography time in fly-scan mode was 13 minutes. Each projection was taken with an exposure time of 350 ms.

### Data analysis

The volume images were loaded in a software for image manipulation and analysis (ImageJ 1.52p by National Institutes of Health, Bethesda, Maryland, USA). Their histograms were cropped to the part of the range with high voxel counts, the bit depth was reduced to 16 bit. The volume images were cropped to the region of interest and loaded in a 3D image analysis software (Amira Software 6.3.0 by Thermo Fisher Scientific Inc., Waltham, Massachusetts, USA). The bone was marked by thresholding and region growing, and the bone surface was extracted. Pairs of landmarks were assigned to common patterns on the bone surfaces of the relaxed state as a reference and of each of the force levels. Based on these landmarks the volume image of each force level was aligned to the reference (Fig. 1B) and subsequently resampled to the reference grid.

The main analysis of deformation was based on visual inspection. For comparison of two co-registered volume images from a specimen, a section plane parallel with the fibers was added. If the orientation of the fibers changed between the volume images, the plane was defined to intersect with three points: a position at the bone surface and a pair of positions that showed correspondance between the force levels. Such pairs of positions should be connected to the position at the bone surface by fibers.

After aligning the section plane, it was used to search for corresponding patterns in three regions along the tendon and to mark them by pairs of landmarks: near the interface with the hard tissue (P0), at the transition from curved fibers in the distal tendon to straight fibers in the free tendon (P1) and in the more proximal free tendon (P2). Position P1 is defined according to the fiber curvature at force level one. The free tendon experienced large displacements between relaxed state and force level one. Therefore, suitable patterns for position P2 were searched for in a large area. Semilandmarks were placed between the landmarks according to the course of the tendon fibers (Fig. 1C,D, 2A–C). We accounted for gross fiber kinks when positioning the semi-landmarks (Fig. 2A), but not for low wavelength crimps as the latter were not resolved in all cases.

The coordinates of landmarks and semilandmarks were exported. A section of the tendon was oriented according to the coordinates of P1 and a normal vector that was calculated from the coordinates of the neighboring semilandmarks. The section was exported to determine tendon CSA. At each force level, the total length (P0–P2), the length of the distal (P0–P1) and the length of the proximal segment (P1–P2) were measured along the course determined by the semilandmarks using a custom software (Cloud2 version 14.11.29 by Heiko Stark, https://starkrats.de). Strains were calculated from the measured lengths:

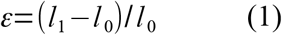

*ε*: strain, *l_0_*: length in the relaxed state, *l_1_*: length at the corresponding force level.

The mean force was calculated from the forces logged during tomography. Stress was calculated from the mean force and the tendon CSA at P1. Tangent moduli of the stress-strain curve were approximated by the slope of the line connecting neighboring data points.

To measure the relative volume change during the force levels, the tendon was segmented in each volume image and exported as a surface. The coordinates of the landmarks P1 and P2 and the neighboring semilandmarks were used to define section planes. The tendon surfaces were cut into separate closed surfaces at these section planes. The volume enclosed by each of the resulting surfaces was measured using a 3D computer graphics software (Blender 2.79b by Blender Foundation, Amsterdam, Netherlands). The relative volume changes were calculated from the volumes in the relaxed state and at each of the force levels.

The volume of the bursa was measured in the same way in two specimens, both unwatered, one measured with an angle of load application of 90°, the other with an angle of 120°.

### Optical flow analysis

In one unwatered specimen, the change in tissue structure between force level one and two was sufficiently small to track tissue displacements automatically. The analysis was performed from dense motion estimates of the tendon, the *Calcaneus* and the surrounding soft tissue. The displacement fields were calculated under the assumption of grey value constancy with a custom global 3D variational optical flow solver implemented in CUDA. Input data were filtered for noise and artifacts with an iterative non-local means filter^[58]^ and global image contrast was enhanced by histogram equalization. This preprocessing routine emphasizes the contrast of textures related to tendon fibers ensuring high quality tracking of deformations in the region of interest. Larger displacements were captured by applying the optical flow algorithm to an image pyramid and using an inner/outer iteration scheme.^[59]^ Best results in preserving flow discontinuities while interpolating untextured and artifact heavy regions were achieved by using anisotropic flow-driven regularization,^[60]^ an L1 penalty term,^[59]^ fourth-order centered finite differences for derivatives and cubic interpolation for warping. Resulting displacement vectors were mapped onto a tetrahedral mesh of the tendon for visualization purposes. The result was rotated for the calculation of the strain tensor to align the tendon orientation with the y-axis of the coordinate system and to place the x-axis in the sagittal plane. The z-axis correspondingly is normal to the sagittal plane. The tensor components of the Green-Lagrange strain tensor were calculated for the elements of the tetrahedral mesh with this coordinate system using a scientific open-source visualization software (ParaView 5.8.0 by Sandia National Laboratories, Kitware Inc. and Los Alamos National Laboratory, all in the USA). The landmarks and semilandmarks that correspond to the relaxed state in the visual analysis were as well displaced using the displacement fields from the optical flow analysis to extract strain values for comparison.

### Statistics

Data from the three specimens in each of the groups (“watered”, “unwatered”) were summarized giving the mean value and the standard deviation. The samples were tested for normal distribution using the test by David et al. (α = 0.05, n = 3). The homogeneity of variances was examined using the F-test (α = 0.05, ν_1_ = ν_2_ = 3). The samples were tested for differences with regard to their central tendencies using Student’s t-test (α = 0.05, ν = 4), Welch’s t-test (α = 0.05) and the U-test (α = 0.05) depending on the type of distribution and homogeneity of variances.

## Acknowledgement

The authors hereby thank Benjamin Naumann, George-Philipp Franz, Agustín Jorge Elias-Costa and David Junghanns who worked with them during the imaging periods at the beamline, Alexander Hipp and Felix Beckmann for knowledgeable support during the assembly and operation of the experiments, Elisabeth Meier and Ms. Brinkmann for keeping the animals, Sabine Bischoff and Michael van der Wall for advice with regard to animal provision and animal rights, Dirk Arnold for recommendations with regard to tissue preparation, Martin S. Fischer, Jörg Bossert, Hartmut Witte, Cornelius Schilling and Christian Rode for the discussion of study design and mechanical interpretation. This study was supported by a grant from Deutsches Elektronen-Synchrotron (DESY: I-20170614) to Martin S. Fischer and Julian Sartori. This research was supported in part through the Maxwell computational resources operated at DESY.

## Supplementary material

Figure S1: Control test without synchrotron radiation.

Figure S2: Stress over strain in individual specimens.

Figure S3: Volume changes in the proximal and distal region in individual specimens.

Figure S4: Relative volume change increases with length of the proximal measuring distance.

Figure S5: Relative volume changes of the complete examined region in the individual specimens.

Figure S6: Relative volume changes of the proximal and the distal part of the examined region.

Figure S7: Dependency of relative volume changes in the proximal part from the initial length of the proximal test distance.

Figure S8: The bursa.

Figure legends are provided with the supplementary figures.

## Table of contents

The article presents the method and the results of the first tomographic deformation study of intact tendon-bone insertions at microscopic resolutions. The method is described and evaluated in detail. Under load, higher strains and volume losses are found to occur in the distal tendon near the insertion than in the more proximal free tendon. This finding confirms previous findings from dissected tendon-bone insertions.

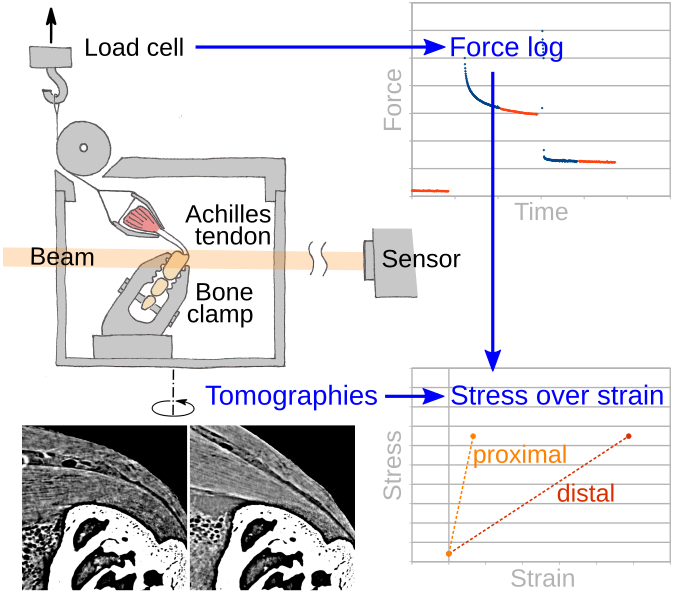

