## Supplemental figures S1-S8 for "Gaining Insight into the Deformation of Achilles Tendon Entheses in Mice"

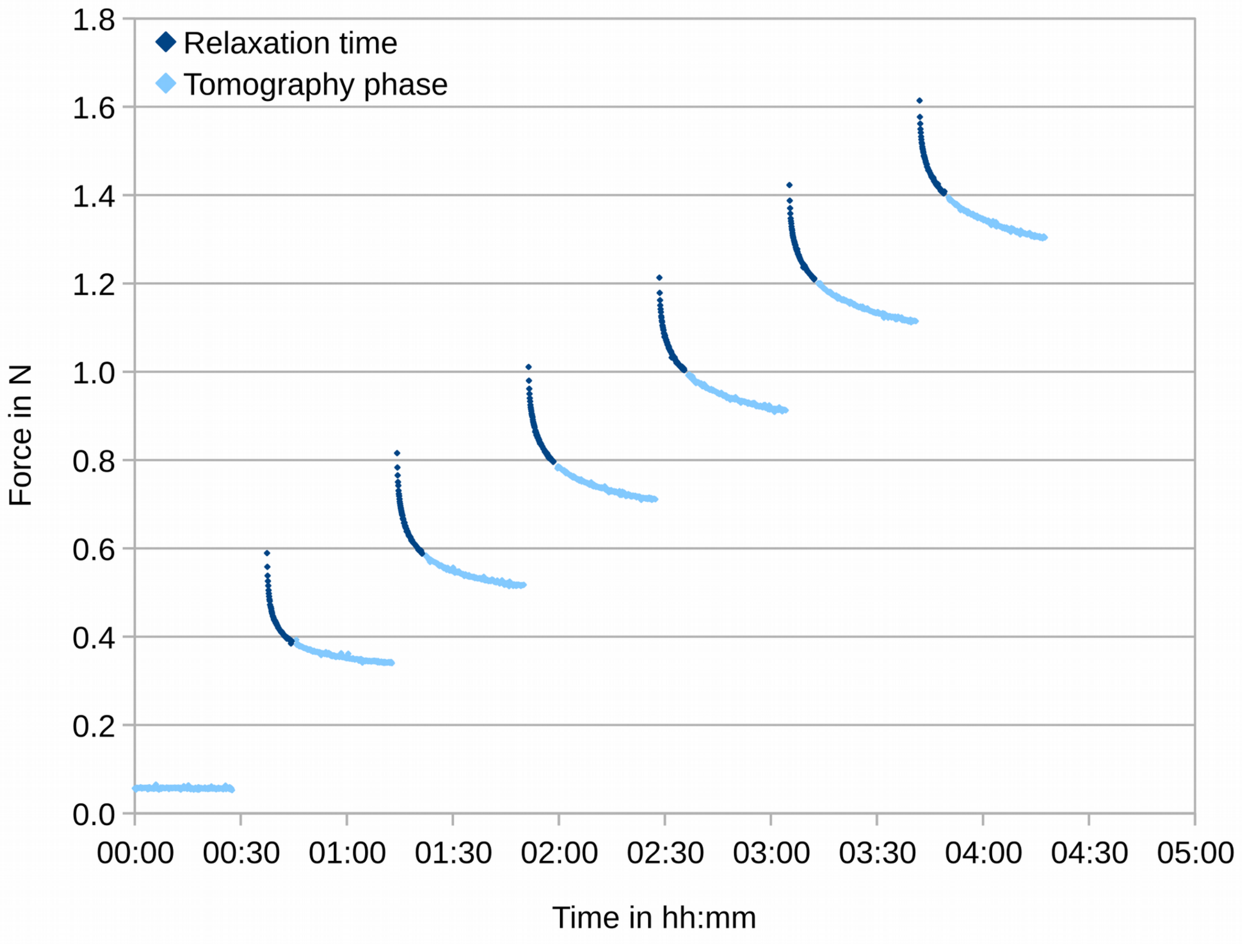


Fig. S1: Force diagram of a specimen tested in a force program without being exposed to synchrotron X-ray radiation during the tomography phases.

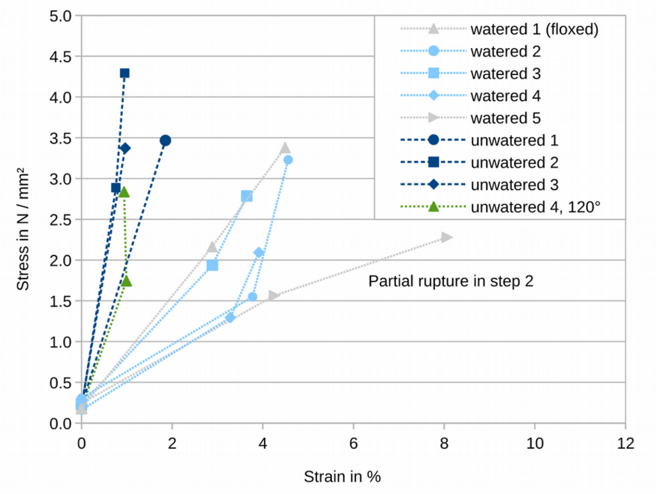


Fig. S2: Stress over strain for the total test distance in the individual specimens. If data from two force steps was available, it is plotted.

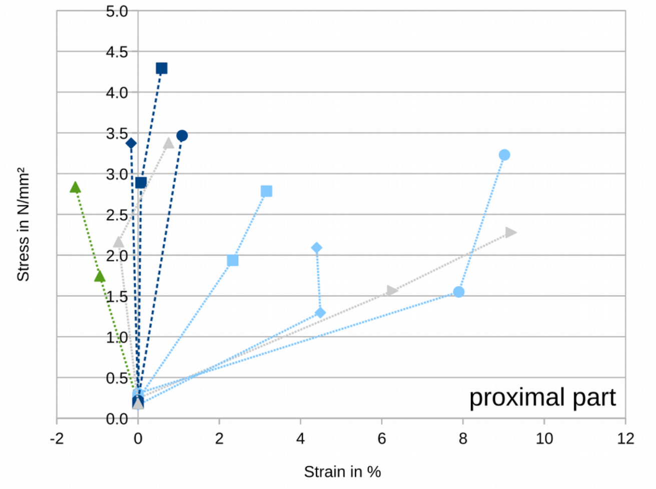


Fig. S3: Stress over strain for the proximal part of the test distance in the individual specimens. If data from two force steps was available, it is plotted. The legend corresponds to figure S2.

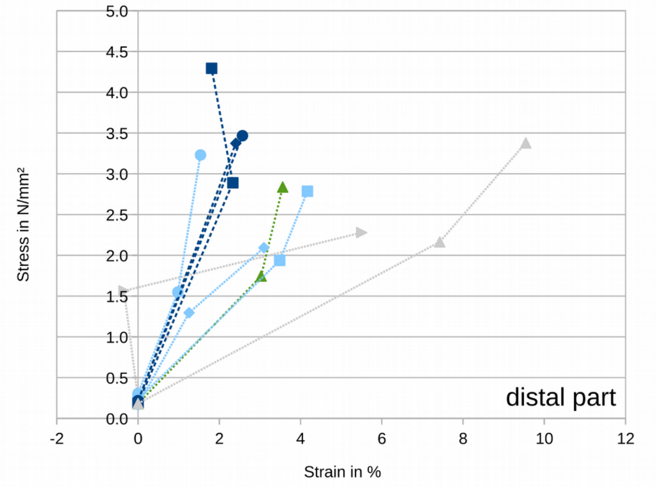


Fig. S4: Stress over strain for the distal part of the test distance in the individual specimens. If data from two force steps was available, it is plotted. The legend corresponds to figure S2.

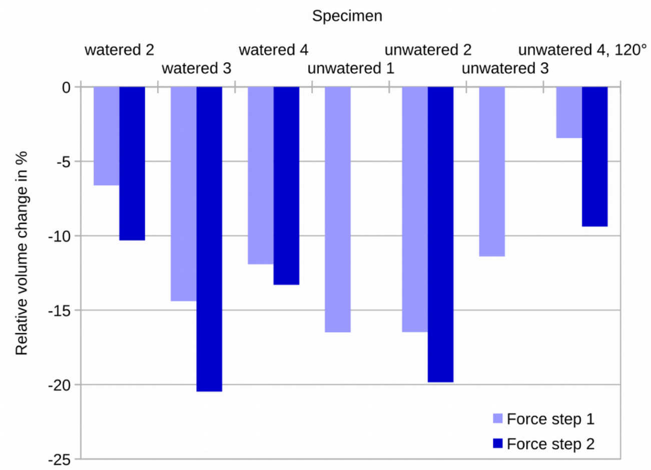


Fig. S5: Relative volume changes of the examined part of the tendon in the individual specimens measured after the first force step and if possible after the second one.

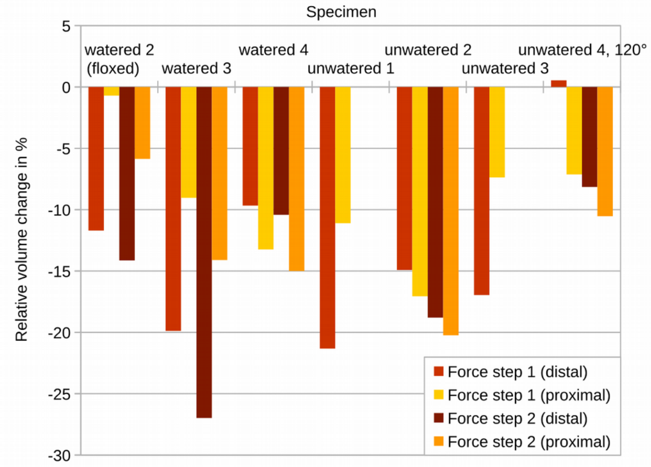


Fig. S6: Relative volume changes of the proximal and distal part of the examined volume in the individual specimens measured after the first force step and if possible after the second one.

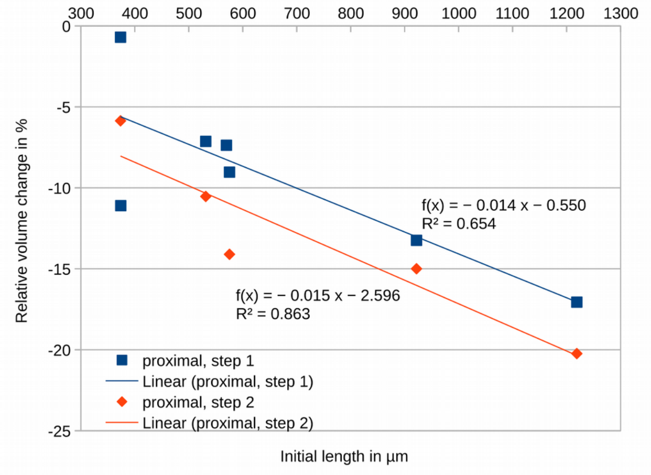


Fig. S7: Relative volume changes in the proximal part over initial length of the proximal test distance. The further the proximal test distance extends into the free tendon, the larger is the decrease in volume in the examined force steps.

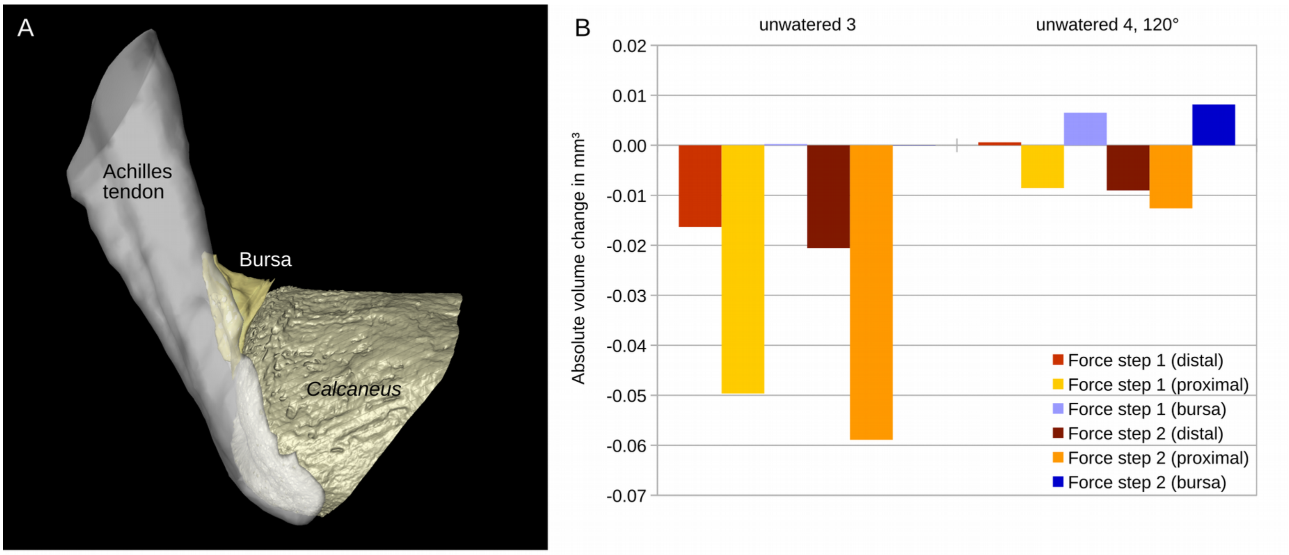


Fig. S8: The bursa. (A) Surface rendering of bone, tendon and bursa. (B) Absolute volume changes of tendon and bursa at nominal insertion angles of 90° and 120°. Each angle is examined in one specimen.
